# Neuromolecular and behavioral effects of ethanol deprivation in *Drosophila*

**DOI:** 10.1101/2021.01.02.425101

**Authors:** Natalie M. D’Silva, Katie S. McCullar, Ashley M. Conard, Tyler Blackwater, Reza Azanchi, Ulrike Heberlein, Erica Larschan, Karla R. Kaun

## Abstract

Alcohol use disorder (AUD) is characterized by loss of control in limiting alcohol intake. This may involve intermittent periods of abstinence followed by alcohol seeking and, consequently, relapse. However, little is understood of the molecular mechanisms underlying the impact of alcohol deprivation on behavior. Using a new *Drosophila melanogaster* repeated intermittent alcohol exposure model, we sought to identify how ethanol deprivation alters spontaneous behavior, determine the associated neural structures, and reveal correlated changes in brain gene expression. We found that repeated intermittent ethanol-odor exposures followed by ethanol-deprivation dynamically induces behaviors associated with a negative affect state. Although behavioral states broadly mapped to many brain regions, persistent changes in social behaviors mapped to the mushroom body and surrounding neuropil. This occurred concurrently with changes in expression of genes associated with sensory responses, neural plasticity, and immunity. Like social behaviors, immune response genes were upregulated following three-day repeated intermittent ethanol-odor exposures and persisted with one or two days of ethanol-deprivation, suggesting an enduring change in molecular function. Our study provides a framework for identifying how ethanol deprivation alters behavior with correlated underlying circuit and molecular changes.

## Introduction

Alcohol use disorder (AUD) is a chronic disorder characterized by, among other behaviors, a loss of control in limiting intake. Individuals suffering from AUD often struggle with recurring periods of abstinence and relapse. Relapse can be provoked by memory of the pleasurable effect of alcohol when faced with alcohol-associated cues [1–6]. In some users, reactivity to alcohol-associated cues has been associated with increases with chronic alcohol use [7–10]. However, the molecular mechanisms underlying formation of alcohol-associated memories is not well understood.

Repeated alcohol use can also result in lasting behavioral adaptations including altered feeding, memory, exploratory behavior, social behavior, increased anxiety and enhanced sensitivity to stress [11, 21, 22]. Alcohol deprivation can exacerbate these behaviors [23–25], which may enhance vulnerability to relapse [26]. Understanding the consequences of neural adaptations on behavior will help us understand the mechanisms that underlie the transition from occasional alcohol use to chronic use, providing a framework for identifying new targets for diagnosis, treatment, and prevention of AUD.

Alcohol alters brain reward systems, which drives alcohol seeking behavior [11, 12]. This includes dysregulation of gene expression, multiple neurotransmitter interactions, and subtle changes in neurochemical function [2, 15–19]. Repeated alcohol use can result in lasting behavioral adaptations including altered feeding, memory, exploratory behavior, social behavior, increased anxiety and enhanced sensitivity to stress [11, 21, 22]. Alcohol deprivation can exacerbate these behaviors [23–25], which may enhance vulnerability to relapse [26]. Understanding the consequences of neural adaptations on behavior will help us understand the mechanisms that underlie the transition from occasional alcohol use to chronic use, providing a framework for identifying new targets for diagnosis, treatment, and prevention of AUD.

*Drosophila* has proven to be a powerful model to elucidate the genetic and molecular underpinnings of alcohol sensitivity, tolerance, and memory [27–31]. *Drosophila* have similar responses to alcohol compared to humans; they actively seek alcohol, become intoxicated, and develop tolerance [29, 32–34]. Recent advancements in computer-vision and machine learning has recently revealed complex interactions between flies that are noticeably modified by changes in the environment [35–40]. *Drosophila* therefore serves as a powerful model to understand how alcohol simultaneously alters social behavior and corresponding gene expression.

Our goal here is to understand molecular and behavioral changes that occur with acute cycles of ethanol-odor exposure and one or two days of forced ethanol deprivation. Although previous studies investigated the behavioral states [41–43] and molecular changes [44, 45] associated with withdrawal, this work uses a three-pronged approach to simultaneously measure and compare behavioral, circuit and gene expression changes. We compare the behavioral and gene-expression shifts that occur following one or three cycles of repeated intermittent ethanol-odor pairings followed by 24 hours of ethanol deprivation. We use high-content tracking data to define spontaneous behavioral shifts, and demonstrate that repeated alcohol exposures alter social behavior for days following exposure. We further identify circuit architecture in the brain correlated with ethanol-modified behaviors using a database of behavior-anatomy maps. We then identify gene expression changes that persist following an additional 24 hours of ethanol deprivation. Together, our results demonstrate that repeated ethanol-odor association cycles induce distinct behavioral states that are associated with specific circuit architecture and gene expression changes. This provides a unique opportunity to form predictions about changes in gene expression and circuit plasticity that coincide with repeated alcohol useabstinence cycles.

## Results

### Ethanol is rewarding to flies after daily intermittent ethanol-odor associations

Memory for alcohol-associated cues has been demonstrated in a wide range of species, from insects to humans [46–52]. It is critical to account for ethanol-cue associations rather than ethanol exposure alone when looking at how ethanol induces neural adaptations, because gene expression changes that occur as a result of alcohol-cue pairings may differ from those that occur when alcohol is presented in the absence of cues [13]. We previously demonstrated that *Drosophila* experience the pharmacological properties of ethanol as rewarding after a single day of three repeated ethanol-odor exposures [11, 21]. How multiple days of this repeated intermittent treatment affects this memory, and how ethanol deprivation might affect this response are unknown.

To first determine if flies experience ethanol intoxication as rewarding after repeated intermittent volatilized ethanol exposure followed by forced ethanol deprivation, we trained wildtype flies to associate ethanol with an odor for one day (Figure 1Ai) or three days (Figure 1Aii, 1Aiii) and measured memory following 24 or 48 hours of ethanol deprivation. Following ethanol-odor training, preference for ethanol was determined by counting the number of flies that move towards an ethanol-paired or ethanol-unpaired odor in a Y-maze (Figure 1B) [11, 21]. Flies showed a significant preference for the ethanol-paired odor 24 hours after three intermittent ethanol-odor pairings (Figure 1Bi), 24 hours after three consecutive days of three intermittent ethanol-odor pairings (Figure 1Bii), and 48 hours after three consecutive days of three intermittent ethanol-odor pairings (Figure 1Biii). Flies exposed to odors alone without ethanol did not show preference for the two odors when tested at the same intervals (Figure S1A-C).

**Figure 1:**
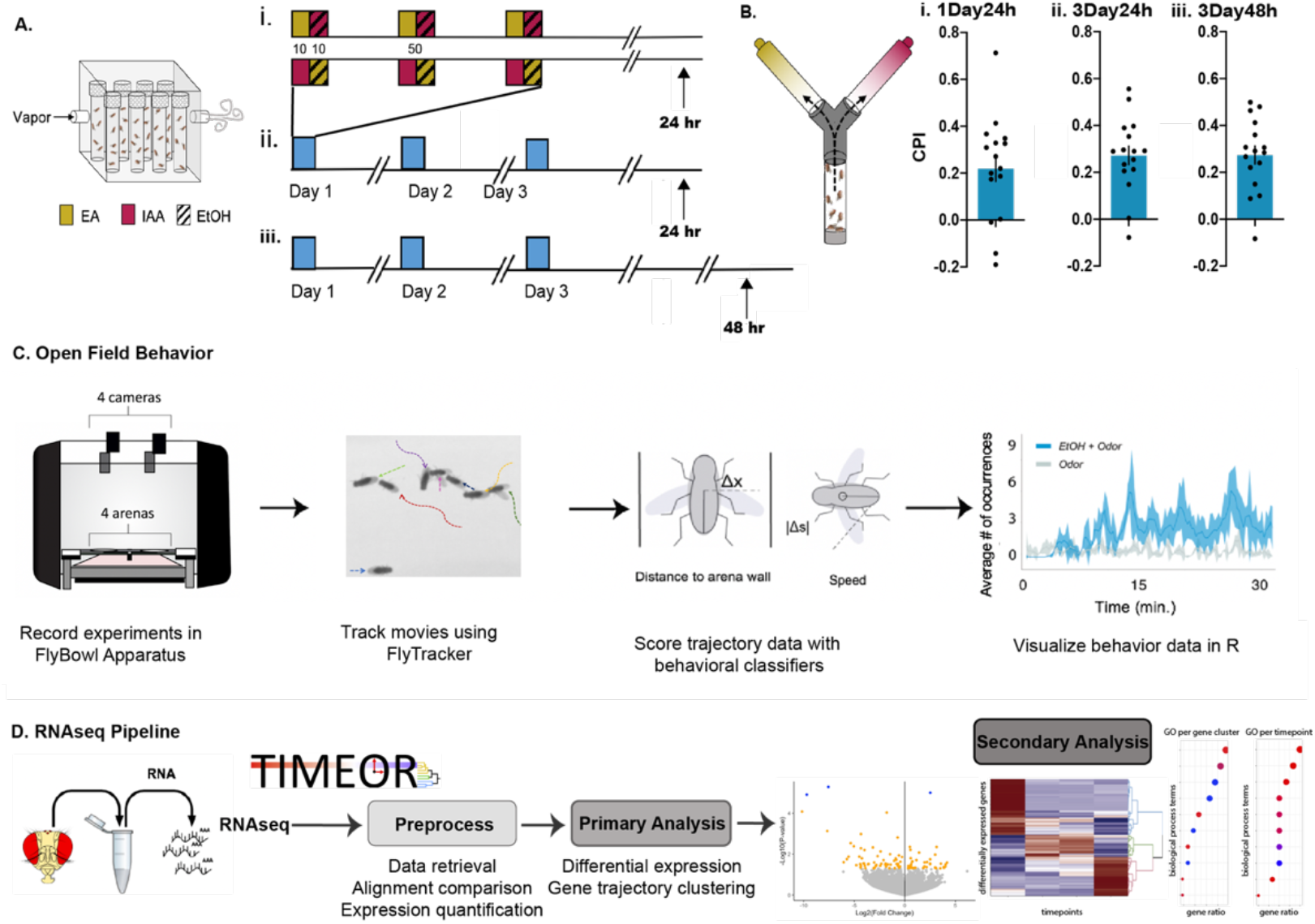
Apparatus and experimental design. (A) Schematic illustrating spaced intermittent alcohol-odor exposure paradigms, for lasting cue-associated alcohol preference. On each day of training, vials of 30 flies are presented with three sessions of 10 min of an unpaired odor, followed by 10 min of paired odor with ethanol. A moderate dose of ethanol (90:60 EtOH:Air) was paired with either isoamyl alcohol or ethyl acetate. Reciprocally trained flies were used to control for odor identity. Flies then underwent forced abstinence for a period of (i) 24 h after one day of spaced intermittent training, (ii) 24 h after three days of spaced intermittent training, or (iii) 48 h after three days of spaced intermittent training. Following forced abstinence, flies were then tested for alcohol-associative memory in a (B) standard Y-maze after (i) 24 h after one day of spaced intermittent training (1Day24h; n = 16), (ii) 24 h after three days of spaced intermittent training (3Day24h; n = 16), or (iii) 48 h after three days of spaced intermittent training (3Day48h; n = 16). (C) Fly behavior was observed in the FlyBowl apparatus, following forced abstinence. The FlyBowl apparatus is an open-field walking arena that accommodates 4 groups of 10 male flies in each of the 4 behavior arenas, in which fly behavior is video recorded. We used automated tracking and behavior analysis algorithms to generate behavioral output data sets from the video recordings. Each behavior (attempted copulation, back up, chaining, chase, crabwalk, jump, pivot center, pivot tail, righting, stops, touch, and walk) was then quantified across time for both the odor control and odor trained groups. (D) Differential gene expression analyses were performed on 30 whole fly heads after the period of forced abstinence. We used an RNAseq processing pipeline, within TIMEOR, to generate and visualize gene expression data. We used R to create volcano plots to display fold change vs p-value statistics between odor control and odor trained libraries.

### Intermittent ethanol-odor exposure alters spontaneous behavior

To test whether one day of intermittent ethanol-odor exposure induced functional changes on behavior, we observed how three intermittent ethanol-odor pairings affected spontaneous behavior in an open field assay 24 hours after the last ethanol-odor pairing (Figure 1C). In order to identify changes in behavior specific to ethanol, we compared animals given ethanol-odor associations to those that receive odor cues in the absence of ethanol (Figure 2A).

**Figure 2:**
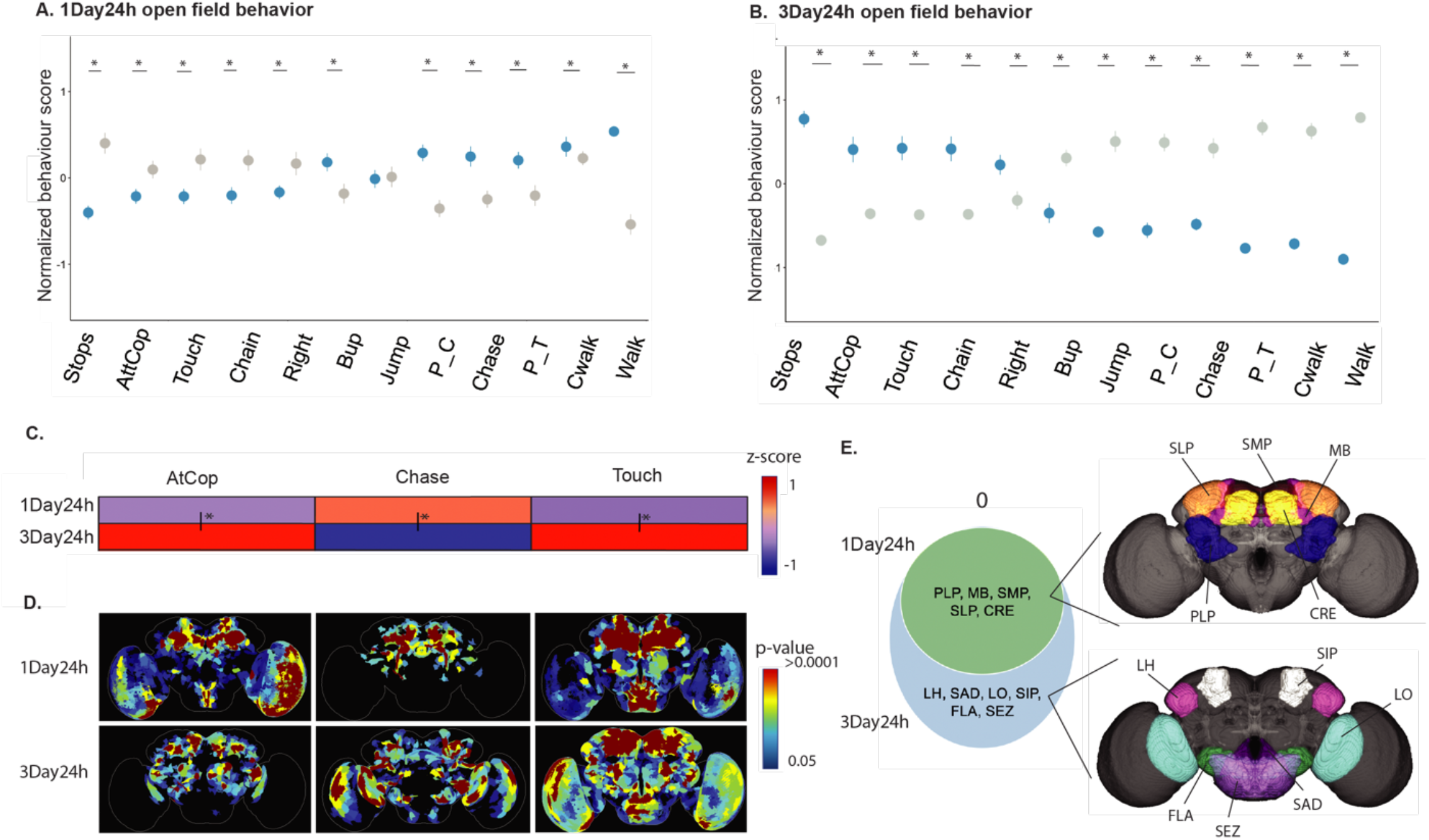
24 hours of forced alcohol abstinence following one day or three day intermittent alcohol paradigms induces behavioral and circuit changes. (A) Open field behavioral trends following 24 hours of forced alcohol abstinence, Mean +/- SEM with statistical significance evaluated using a Student’s T-Test, *p<0.05 following one day of intermittent alcohol exposure (1Day24h; n=8,8), (B) following three days of intermittent alcohol exposure (3Day24h; n=7,8). (C) 1Day24h and 3Day24h EtOH group behavior occurrences sums compared to the odor controls total sums. Red indicates an increase in total sum and blue indicates a decrease in total sum. Statistical significance was evaluated by ANOVA, *post hoc* Tukey, to compare EtOH trends across conditions, *p<0.05. Below the heatmap, we show (D) BABAM projections of the brain regions implicated in the behavioral trends found for each behavioral feature. Dark red indicated the lowest p-value, blue indicates the highest p-value. (E) A Venn diagram showing the brain regions implicated in the social behavioral trends found in each condition. No brain regions were found unique to 1Day24h of ethanol-odor exposure. The PLP, MB, SMP, SLP, and CRE were common to both 1Day24h and 3Day24h of ethanol-odor exposure. The LH, SAD, LO, SIP, FLA, SEZ were found to be unique to the three days of exposure social behavior trends. The brain regions in each section of the Venn diagram are projected onto the brain maps to the right of the Venn diagram.

Freely behaving groups of 10 flies were observed in an open-field walking arena, called the FlyBowl for 30 minutes [53]. Behavioral features of each fly were measured using the computer vision tracking software FlyTracker [54] and classified using the machine learning behavioral classifier JAABA [55] (Figure S2). If a classified behavior was able to be performed by a single fly, the behavior was categorized as a locomotor behavior. This included stopping (Stop), backing up (Bup), righting itself to an upright posture (Right), jumping (Jump), turning (pivot center: P_C or pivot tail: P_T), walking (Walk) and walking sideways (crabwalk or Cwalk). Any behavior requiring two or more flies to interact, was categorized as a social behavior. Social behaviors included touching (Touch), chasing (Chase), attempting copulation (AtCop), and chaining (Chain), in which male flies will form a chain of at least 3 flies with each fly courting the fly in front of it.

Compared to odor-only controls, flies that received ethanol-odor pairings show a significant increase in locomotor behaviors including backing up (p=0.02), crabwalking (p<0.0001), turning (pivot center p<0.0001, pivot tail p=0.009), and walking (p<0.0001) (Figure 2A, S2B, S2E, S2G, S2H, S2L; Data S1). Similarly, they showed a significant decrease in righting (p=0.04) and stopping (p<0.0001) (Figure 2A, S2I, S2J; Data S1). The flies that received ethanol-odor pairings also showed a significant change in social behaviors including a decrease in attempted copulation (p=0.007), chaining (p=0.01), and touching (p=0.0006), and a significant increase in chasing (p=0.002) (Figure 2A, S2A, S2C, S2D, S2K). No significant changes were noted in jumping (p=0.9) (Figure 2A, S2F; Data S1). Thus, for the most part locomotor behaviors increase and social behaviors decrease after three spaced ethanol-odor pairings followed by 24 hours of ethanol-deprivation.

### Repeated intermittent ethanol-odor exposure alters spontaneous behavior

We hypothesized that flies that receive multiple days of spaced ethanol-odor pairings followed by a 24 hour period of ethanol-deprivation would demonstrate a marked shift in locomotor and social behaviors. Flies received three intermittent ethanol-odor pairings for three consecutive days and spontaneous behavior in the FlyBowl was tested 24 hours later (Figure 1Aii, S3). Compared to odor-only controls, flies receiving ethanol-odor pairings showed reduced backing up (p<0.0001), chasing (p<0.0001), crabwalking (p<0.0001), jumping (p<0.0001), turning (pivot center p<0.0001, pivot tail p<0.0001), stopping (p<0.0001), and walking (p<0.0001) (Figure 2B; Supplemental Fig 3B, 3E-J, 3L; Data S1). They also increased attempted copulations (p<0.0001), chaining (p<0.0001), and touching (p<0.0001) (Figure 2B, S3A, S3C, S3D, S3K: Data S1). Thus, locomotor behaviors decrease and social behaviors increase following three consecutive days of three intermittent ethanol-odor pairings.

To investigate if behavioral trends change with additional days of ethanol-odor training, we normalized ethanol-odor treatment to odor only treatment for each behavior, then compared one day ethanol-odor training to three days of ethanol-odor training (Figure 2C, S4). We found significant shifts in all behaviors between the two ethanol-odor training paradigms except pivot tail and backup (Figure S3). There was a pronounced reversal in social behaviors: touch and attempted copulation decreased after one day of ethanol-odor training, but increased after three days of ethanol-odor training, whereas chasing increased after one day of ethanol-odor training but decreased after three days of ethanol-odor training (Figure 2C).

### Repeated intermittent ethanol-odor exposure alters neural circuits

The marked change in open field behavior associated with increased ethanol-odor associations suggests that neuroadaptations are occurring within the circuits mediating these behaviors. To identify circuits associated with changes in behaviors, we took advantage of the Browsable Atlas of Behavior-Anatomy Maps (BABAM) software, which contains information for all behavior classifiers associated with activation of 2,204 genetically targeted neuronal populations [36]. After determining that a behavioral classifier from our dataset was significantly different in the ethanol-odor paired group, this classifier, and the direction of change (increased or decreased in comparison to control) was inputted into the BABAM software and a brain projection produced with the brain regions that are associated with this shift in behavior.

Brain regions associated with social behaviors that were significantly different in the ethanol-odor paired group after one day of ethanol-odor cycles include the posterior lateral protocerebrum (PLP), mushroom body (MB), superior medial protocerebrum (SMP), superior lateral protocerebrum (SLP), and the crepine (CRE) (Figure 2D, 2E; Data S2). Brain regions associated with locomotor behavioral shifts that were changed 24 hours after one day of ethanol-odor associations include the medulla (ME) (Figure S4; Data S2).

Brain regions associated with social behaviors that were significantly different in the ethanol-odor paired group after three days of ethanol-odor cycles similarly include the protocerebrum (PLP), mushroom body (MB), superior lateral protocerebrum (SMP), superior lateral protocerebrum (SLP), crepine (CRE), and the anterior ventrolateral protocerebrum (AVLP), but also include the lateral horn (LH), lobula (LO), superior intermediate protocerebrum (SIP), flange (FLA), saddle (SAD), and subesophageal zone (SEZ) (Figure 2D, 2E; Data S3). Brain regions associated with locomotor behavioral shifts 24 hours following three days of ethanol-odor associations include the posterior lateral protocerebrum (PLP) and the subesophageal zone (SEZ) (Figure S4; Data S3).

### Intermittent ethanol-odor exposure alters transcription

To understand the molecular changes that occur with the pronounced behavioral shifts, we investigated transcriptional changes in the heads of flies receiving ethanol-odor associations compared to age-matched sibling control flies that received odor in the absence of ethanol (Figure 1D). Following one day of ethanol-odor pairings, the expression of *CG34212, CG16826* were downregulated, and *Cyp4g1* was upregulated, using a conservative adjusted p-value of <0.1 (Figure 3A, Data S4). Gene ontology (GO) analysis using a less conservative non-adjusted p<0.05 cut-off revealed responses related to olfaction including “response to pheromone”, “sensory perception of chemical stimulus”, “detection of pheromone”, and “sensory perception” (Figure 3B, Data S5).

**Figure 3:**
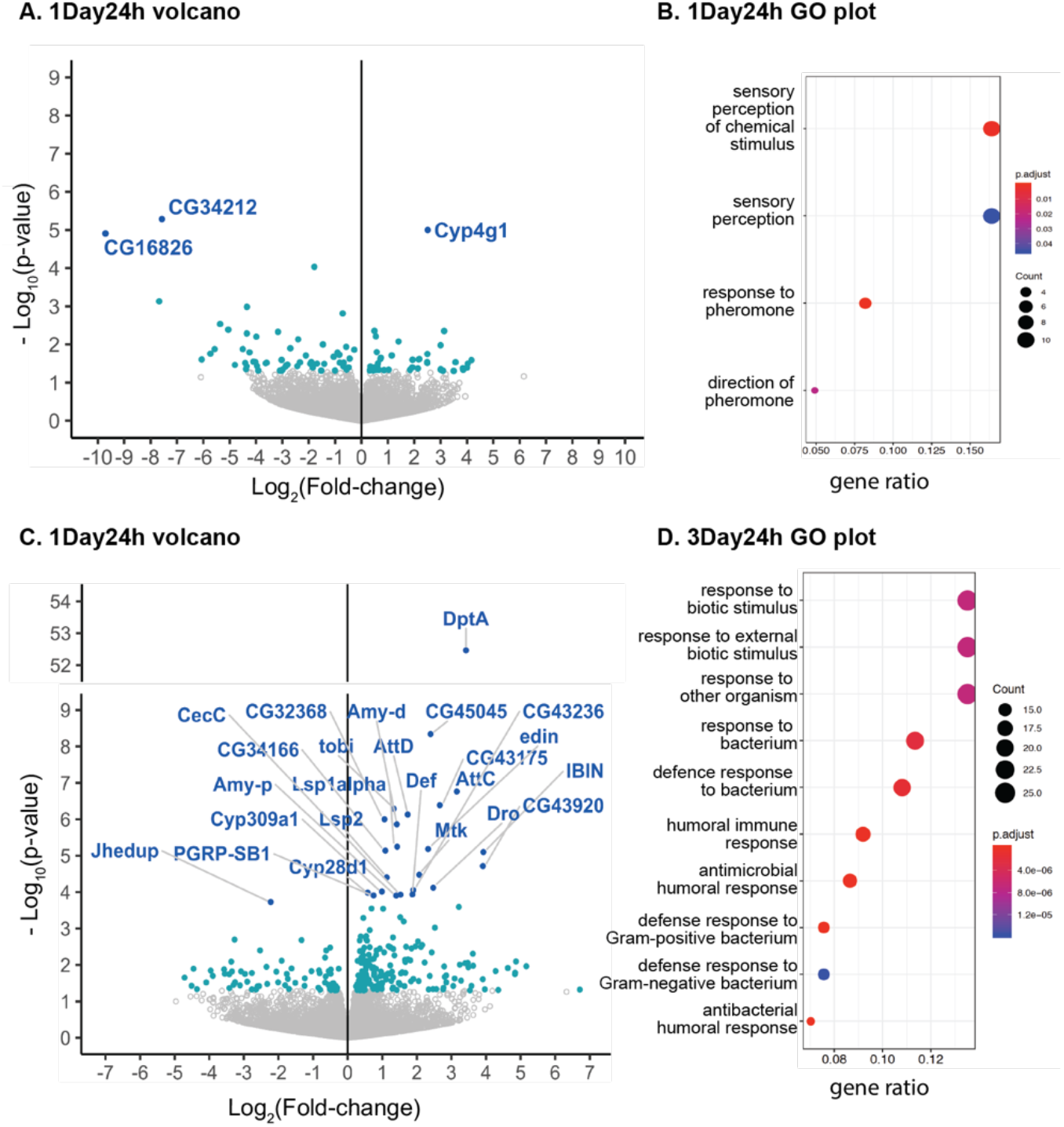
24 hours of forced alcohol abstinence following one day or three day intermittent alcohol paradigms induces gene expression changes. Differential gene expression analyses were performed 24 h after one day of spaced intermittent training (1Day24h) (A) The volcano plot displays fold change *vs* p-value statistics between odor control and odor trained libraries, showing p-value < 0.05 (blue-green), and adjusted p-value of <0.1 (blue; genes labelled). (B) Enriched GO terms for genes with a p-value < 0.05 identified in 1Day24h were plotted on a dot plot. The four GO processes with the largest gene ratios are plotted in order of gene ratio. The size of the dots represents the number of genes associated with the GO term, and the color of the dot represents the p-adjusted values. Differential gene expression analyses were performed 24 h after three days of spaced intermittent training (3Day24h) (C) The volcano plot displays fold change vs p-value statistics between odor control and odor trained libraries, showing p-value < 0.05 (blue-green), and adjusted p-value of <0.1 (blue; genes labelled). (D) Enriched GO terms for genes with a p-value < 0.05 identified in 3Day24h were plotted on a dot plot. The 10 GO processes with the largest gene ratios are plotted in order of gene ratio. The size of the dots represents the number of genes associated with the GO term, and the color of the dot represents the p-adjusted values.

Following three days of ethanol-odor pairings, 24 genes were differentially regulated using a conservative adjusted p-value of < 0.1. *Jhedup* was downregulated, and *DptA, CR45045, AttC, CG43175, tobi, AttD, CG32368, Amy-d, Lsp1alpha, edin, CG34166, IBIN, CG43920, Mtk, Lsp2, Dro, CG43236, Cyp309a1, Amy-p, CecC, Def, Cyp28d1, PGRP-SB1* were upregulated (Figure 3C, Data S5). GO enrichment analysis on differentially expressed genes with a less conservative non-adjusted p<0.05 cut-off revealed terms related to immunity including “response to biotic stimulus”, “defense response to bacterium”, “humoral and antimicrobial immune response” (Figure 3D, Data S5). Since the only difference in treatment between experimental and control flies was the exposure to alcohol, we believe gene expression changes are specific to alcohol treatment rather than age or other environmental experiences.

### Persistent effects of repeated intermittent ethanol-odor exposure

To test whether behavioral state and gene expression changes observed after three days of intermittent alcohol exposure persist, we measured open field behavior and gene expression following an additional 24 hour of ethanol deprivation. We hypothesized that the additional ethanol deprivation might exacerbate the changes in behavior and gene expression. Flies that received repeated intermittent ethanol-odor followed by 48 hours of ethanol deprivation showed a significant increase in attempted copulation (p=0.03), backing up (p=0.004), chaining (p=0.003), righting (p=0.004), and touching (p<0.0001), and a significant decrease in stopping (p<0.0001) (Figure 4A, S5; Data S1). The increase in multiple social behaviors including attempted copulation, chaining and touching, was largely consistent with changes seen after 24 hours of ethanol deprivation, whereas most locomotor behaviors no longer showed significant difference between ethanol-odor and odor-only treated flies. This suggests persistent changes in multiple social behaviors following 48 hours of ethanol deprivation (Figure 4B).

**Figure 4:**
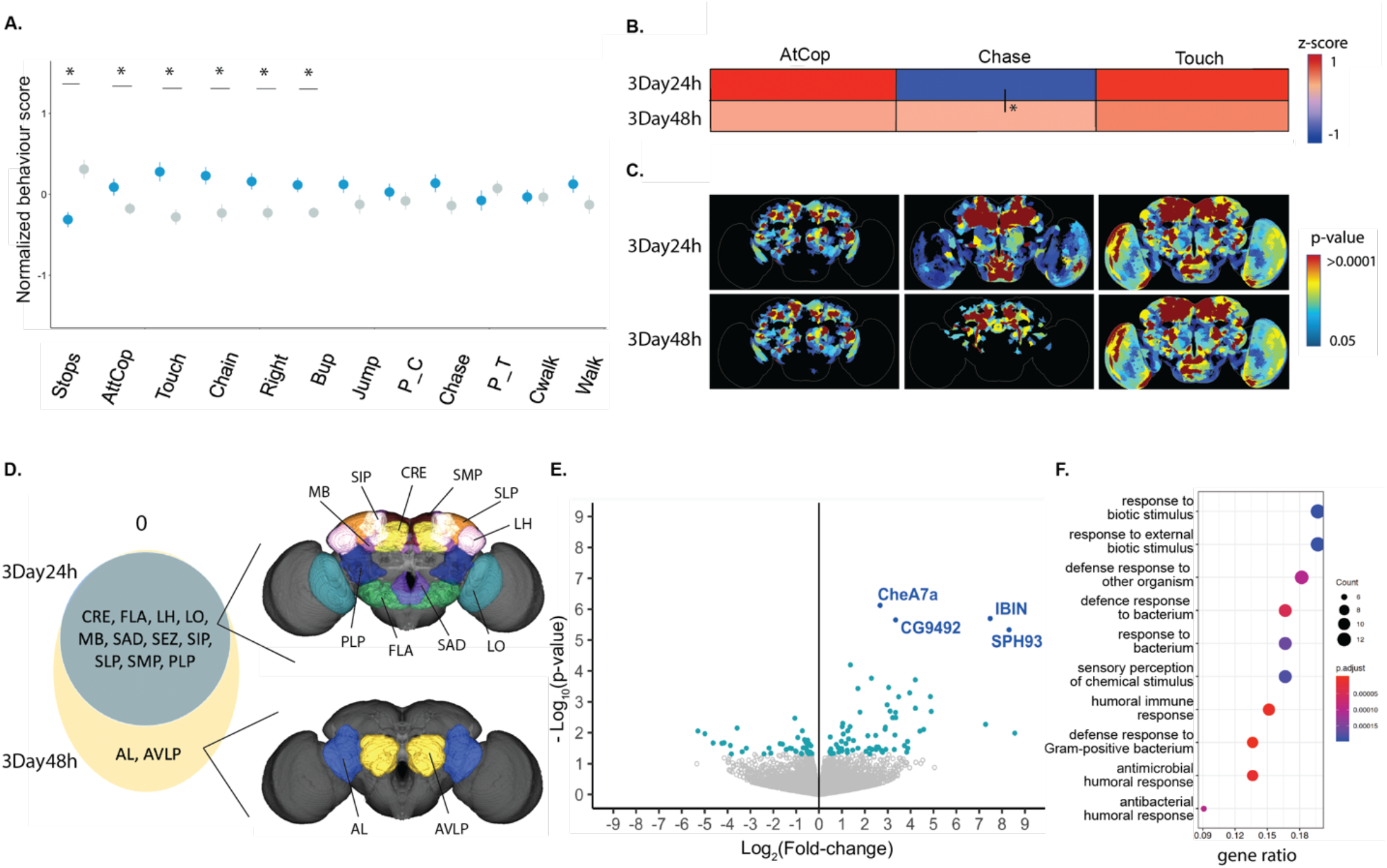
Three days of the intermittent alcohol paradigm induces lasting behavioral, circuit and gene expression changes. (A) Open field behavioral trends following 48 hours of forced alcohol abstinence after 3 days of intermittent alcohol exposure (3Day48h; n=8,8), Mean +/- SEM with statistical significance evaluated using a Student’s T-Test, *p<0.05. (B) Heatmap of the EtOH group behavior occurrences sums compared to the air control total sums at 3Day24h and 3Day48h. Red indicates an increase in total sum and blue indicates a decrease in total sum. Statistical significance was evaluated by ANOVA, *post hoc* Tukey, to compare EtOH trends across conditions, *p<0.05. (C) BABAM projections of the brain regions implicated in the behavioral trends found for each behavioral feature. Dark red indicated the lowest p-value, blue indicates the highest p-value. (D) A Venn diagram showing the brain regions implicated in the social behavioral trends found in each condition. No regions were uniquely identified at 3Day24h. The PLP, MB, SMP, SLP, and CRE were common to both 3Day24h and 3Day48h. The AL and AVLP were implicated only in the social behavior trends seen at 3Day48h. The brain regions in each group are projected onto the brain maps to the right. Differential gene expression analyses were performed at 3Day48h. (E) The volcano plot displays fold change vs p-value statistics between odor control and odor trained libraries, showing p-value < 0.05 (bluegreen), and adjusted p-value of <0.1 (blue; genes labelled). (F) Enriched GO terms for genes with a p-value < 0.05 identified at 3Day48h were plotted on a dot plot. The 10 GO processes with the largest gene ratios are plotted in order of gene ratio. The size of the dots represents the number of genes associated with the GO term, and the color of the dot represents the p-adjusted values.

Brain regions associated with the persistent social behavioral shifts included the mushroom body (MB), superior medial protocerebrum (SMP), superior lateral protocerebrum (SLP), crepine (CRE), anterior ventrolateral protocerebrum (AVLP), lateral horn (LH), antennal lobe (AL), lobula (LO), superior intermediate protocerebrum (SIP), flange (FLA), saddle (SAD), and the subesophageal zone (SEZ) (Figure 4C, 4D; Data S6). The antennal lobe (AL) and the anterior ventrolateral protocerebrum (AVLP) were the only brain regions uniquely associated with the lasting social behavioral changes (Figure 4C, 4D).

In addition to the persistent behavioral changes and circuits associated with 48 hours ethanol deprivation following three days of intermittent alcohol exposure, we also observed persistent immune gene expression changes. Four unique genes, *CheA7a*, *CG9492*, *IBIN*, and *SPH93*, were significantly upregulated with conservative adjusted p-value <0.1 (Figure 4E, Data S7). However, gene ontology analysis of differentially expressed genes selected using a less conservative non-adjusted p<0.05 revealed biological process terms similar to those flies tested 24 hours previously (Figure 3D, 4F). Biological process terms were predominantly associated with an immune response, for example, “defense response to bacterium”, “immune response”, “defense response to other organism”, and “defense response to Gram-positive bacterium” (Figure 4F, Data S8).

### Gene expression changes with ethanol-odor exposure and deprivation

The similarity in GO terms identified after 24 hours or 48 hours of ethanol deprivation following three days of intermittent ethanol-odor training suggested that repeated ethanol exposure induces long-lasting gene expression changes. To highlight the persistent gene expression trajectories, we used the unbiased time-series RNA-seq and multi-omics analysis method Trajectory Inference and Mechanism Exploration with Omics data in R (TIMEOR) to identify differentially expressed genes and how they temporally co-regulate across all conditions, forming gene expression trajectory clusters [56]. TIMEOR compiled a list of differentially expressed genes (using the less conservative p<0.05 cut-off) from each condition (Data S9). TIMEOR then automatically clustered (using unsupervised clustering) the resulting 430 differentially expressed genes into nine groups representing similar gene dynamic trajectories, shown in an interactive clustermap [56] [57] (Figure 5A, S6, Data S10).

**Figure 5:**
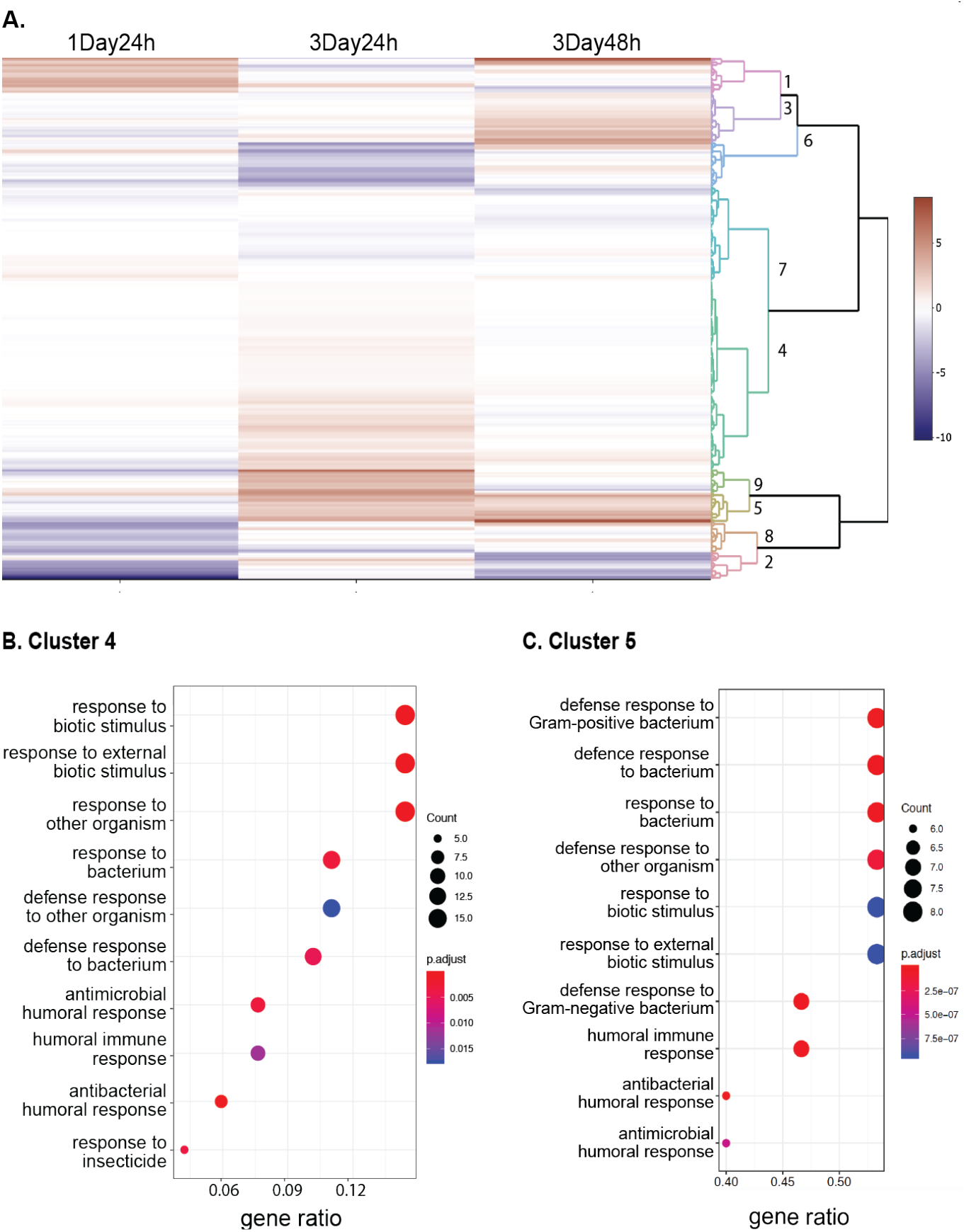
Gene expression changes with alcohol exposure and forced abstinence. (A) We used a clustermap to identify clustering of differentially expressed genes, with a p-value < 0.05, from each condition, 1Day24h, 3Day24h, and 3Day48h, resulting in 9 clusters representing similar gene dynamic trajectories. (B) GO enrichment analysis on genes within cluster 4, which includes genes that increased expression at 3Day24h, were plotted on a dot plot. The ten GO processes with the largest gene ratios are plotted in order of gene ratio. The size of the dots represents the number of genes associated with the GO term, and the color of the dot represents the p-adjusted values. (C) GO enrichment analysis on genes within cluster 5, which includes genes that increased expression at 3Day24h, and maintained increased expression at 3Day48h, resulted in a list of ten GO processes, that were plotted on a dot plot (similar to panel B).

Clusters 3, 4 and 5 had significantly enriched gene ontology (GO) terms. Genes in Cluster 3 were predominantly associated with biological terms related to chemosensory functions, including upregulation of *Obp19a*, *Obp59a*, *Or42b* and *Orco*, 24 hours after one-day of intermittent alcohol exposure (Figure S6C, S6D). In contrast, genes in clusters 4 and 5 contained genes that were persistently changed following three days of intermittent ethanol-odor exposure, and were largely associated with biological process terms predominantly associated with immune response (Figure 5B, 5C). Cluster 4 includes upregulation of *Tsf1*, *aay*, *AttD*, *AttB*, *PGRP-SB1*, and *Tdc1*, at both 24 hours and 48 hours following three days of intermittent ethanol-odor exposure (Figure S6E). Cluster 5 includes upregulation of *CecA1*, *CecA2*, *DptA*, *Dro*, *Mtk*, *DptB* (Figure S6F).

To identify potential connections between genes that emerged from clusters 4 and 5, we used the MIST database [58] and the open source Cytoscape platform [59] to visualize the protein-protein interaction (PPI) network of differentially expressed genes in these clusters (Figure 6A, Data S11, S12). Notably, *Relish* (*Rel*) the primary transcription factor in Immunodeficiency (IMD) pathway, had the largest known network, including a number of other proteins associated with the IMD pathway (*PGRP-LE*, *Fadd*, *Tab2*, *Tak1*, *Dredd*, *PGRP-LC*, *Stat92E*), and nuclear factor-κB signaling complex (*Dif*, *dl*, *cact*). Together, the differentially regulated immune response genes in Clusters 4 and 5 implicate the innate immune signaling pathways Toll and IMD as remaining upregulated as a result of repeated intermittent ethanol-odor pairings followed by either 24 or 48 hours of ethanol deprivation (Figure 6B).

**Figure 6:**
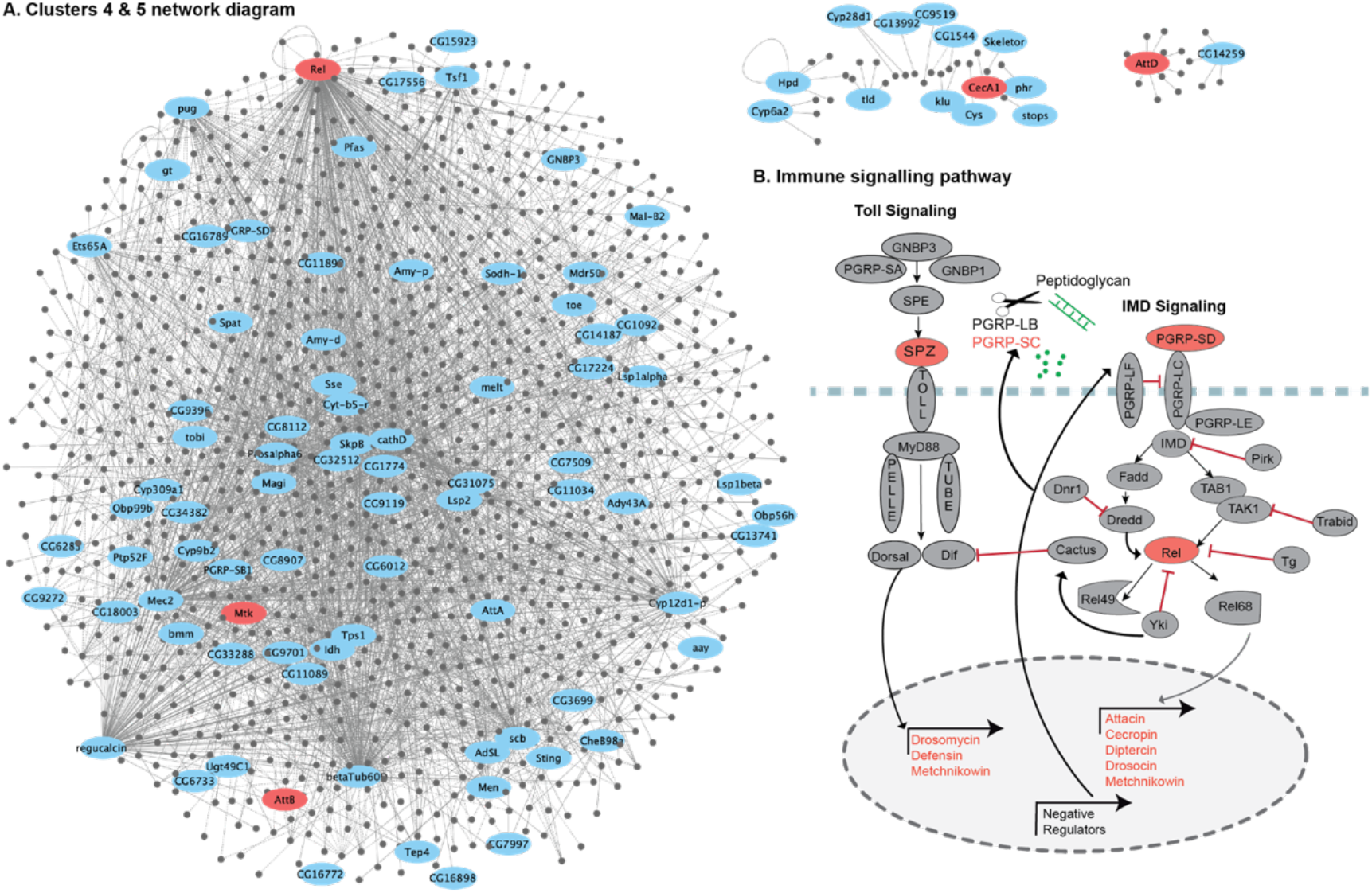
Immune pathway genes upregulated after three days of spaced intermittent training. (A) Protein-protein interaction (PPI) network analysis of genes in clusters 4 and 5. Each node represents a protein, blue ellipses represent differentially expressed genes within cluster 4 or 5, and grey circles represent interactors. Red ellipses represent immune response genes that increased expression at 3Day24h, and maintained increased expression at 3Day48h. Interactions between proteins are denoted by grey lines. Bold lines indicate experimentally identified physical interactions PPI, and dashed lines indicate mapped PPI from other species. (B) Immune response genes identified in panel A are highlighted (red) in a schematic representation of the *Drosophila* Toll and IMD pathways. Differentially expressed immune response genes from all clusters, that increased expression at 3Day24h, and maintained increased expression at 3Day48h, are also highlighted in red within the schematic.

## Discussion

Our study sought to uniquely compare behavioral, neural circuit and molecular changes associated with one or three days of repeated intermittent ethanol-odor pairings followed by ethanol-deprivation. We demonstrate one day of ethanol-odor associations followed by 24-hours of deprivation resulted in increased locomotor and decreased social behaviors, whereas three cycles of the ethanol-odor pairing/deprivation paradigm resulted in decreased locomotor and increased social behaviors. The profound behavioral switch from one to three cycles is associated with eight unique brain structures, and a sustained increase in genes associated with innate immunity. Using our three-pronged approach we establish a framework to measure and compare behavioral, circuit and gene expression changes, thus providing a unique opportunity to identify changes in gene expression that coincide with repeated alcohol use-abstinence cycles, and form predictions on where these changes occur in the brain.

### Alcohol alters behavior state

Elucidating how alcohol alters behavioral state is complicated and multifaceted. Internal motivational state, previous experiences as well as environmental conditions can impact an animal’s behavior [35]. Our data here demonstrate that repeated intermittent ethanol-odor exposure induces an enduring appetitive response to the pharmacological properties of ethanol. Moreover, 24 to 48 hours of ethanol deprivation following repeated intermittent ethanol-odor exposure profoundly affected locomotion and social behavior, suggesting that removal of the reward alters behavioral state.

Following one day of exposure, flies showed increased bouts of locomotion including walking and turning, and decreased social behaviors including touching, attempted copulation and chaining (Figure 2A). However, after three days of exposure, flies showed a reversal in these behavioral trends with decreased bouts of locomotion including walking, jumping and turning and increased social behaviors including touching, attempted copulation and chaining (Figure 2B). Previous studies demonstrate that flies decrease locomotion and increase aggressive social interactions in response to stressful environments [60, 61]. Similar anxiety-induced responses have also been detailed in mammals, commonly referred to as “fight-or-flight” responses characterized by freezing in place, hiding if possible, and engaging in more frequent aggressive social behavior [62, 63]. Similarly, anxiety and aggression in humans are interconnected and can manifest in a fight-or-flight response [64]. We found that changes in *Drosophila* social behavior persist with additional ethanol deprivation, suggesting molecular adaptations induced by ethanol endure within these circuits and are expressed as a negative affect state occurring concurrently with removal of a rewarding stimulus.

### Alcohol may engage the mushroom body to affect social behavior

The focus on ethanol-odor associations in our paradigm predictably indicated the involvement of the mushroom body (MB), a central brain structure that plays a role in learning, memory, attention, feature abstraction, and multisensory integration [65–67]. MB circuits are required for formation of alcohol associated memories [14, 21], and ethanol-odor exposure affects gene expression in the MB [13, 31]. We identified changes in specific locomotor and social behaviors after both one or three days of ethanol-odor pairing that were computationally associated with activity in the MB and the surrounding neuropil including the superior medial protocerebrum (SMP), superior lateral protocerebrum (SLP), and crepine (CRE). We speculate that ethanol-induced changes in MB activity influence behavioral state, including how flies interact with each other.

The MB has also been implicated in complex behavior across many insect species [68]. In honeybees, the MBs are critical for social behavior and experience [69–73]. Valence of social information is encoded in different subpopulations of intrinsic MB neurons [74]. These changes occur as a result of molecular adaptations, since transcriptomic responses in the mushroom body of honeybees differs after social experiences [75, 76]. Similarly, in primitively eusocial bees, the MB is larger among queens which have more complex social roles than workers [77]. Social wasps have larger MBs than solitary wasps [78, 79], and social experience alters MB volume [80, 81]. And finally, socially complex ant species have larger relative investment in the MB than more socially basic species [82]. Similarly, ants of the same species raised in social isolation demonstrate impairment in the growth of the MB compared to ants raised in social groups [83].

In *Drosophila*, the activity of MB circuitry is required for memories for olfactory cues associated with losing fights, demonstrating its requirement for establishing dominant-subordinate male-male relationships [84, 85]. The MB is also required for social transference of memory for the presence of a predator [86]. More recently, social cues encoded in MB gamma neurons were shown to be required for social attraction [87]. Our results add to these findings by implicating the MB and surrounding neuropil in long-lasting adaptations in social behavior by ethanol.

### Alcohol broadly engages sensory regions of the brain

The collective behaviors that emerge as a result of ethanol-odor exposure also engage widespread areas of the brain outside of the MB (Figure 2E, 4D, S3). The persistent changes in social behavior revealed a shift from involvement of the MB and surrounding neuropil to also include the anterior ventrolateral protocerebrum (AVLP), lateral horn (LH), antennal lobe (AL), prow, lobula, superior intermediate protocerebrum (SIP), flange and subesophageal zone (SEZ) (Figure 4D). These brain regions are all involved in sensory detection or integration and are characterized by widespread interconnectivity. This suggests that brain areas required for sensory integration become recruited into neuroadaptive responses initiated as the effects of ethanol deprivation become more prominent.

For example, our data indicates that 24 hrs of ethanol deprivation following three days of intermittent ethanol-odor exposure alters social behavior in a way that engages the entire olfactory system including the first order olfactory neurons (AL) and area primarily responsible for innate responses (LH) in addition to the MB [88–91]. Given the extensive crosstalk between the MB and the LH, it’s likely that ethanol may influence how learned behaviors influence innate responses [92–95]. Similarly, the SIP receives projections from the MB, and contains neurons that project to the central complex [96, 97]. This suggests that ethanol may influence how stimuli and motivated response are integrated before they are relayed to an area with a pronounced role in navigation [98]. Finally, the recruitment of the SEZ is of interest as this region has been previously implicated in social behaviors including courtship and aggression as well sensory processing and locomotor output [99–105]. The broad engagement of sensory detection and integration after ethanol-odor exposure and acute deprivation indicates that ethanol impacts the ability to taste and smell, leading to modified social and locomotion behaviors.

In contrast, social behavior changes that occur after three ethanol-odor pairing cycles and persist following 48 hours of ethanol-deprivation were specifically associated with activity in the medulla (ME). The ME is located in the fly’s optic lobe and processes incoming visual information [106, 107]. Perhaps the involvement of the ME is a result of resetting behavioral strategies from that of olfaction-based to visual-based following alcohol cessation. Taken together this data suggests that alcohol-induced neuroadaptations occur in sensory processing areas, and shift as behavioral strategies change to facilitate navigation in social environments.

### Alcohol alters gene expression associated with sensory response

The observed changes in gene expression with repeated intermittent ethanol-odor exposure are also consistent with a neuroadaptive change in sensory pathways. Genes associated with sensory perception and pheromone response were enriched following one day of intermittent ethanol-odor pairings (Figure 3B). The marked upregulation in a number of odorant binding proteins (OBPs) genes following ethanol-odor pairings appears to be at the intersection of alcohol-induced behavior, social behavior and memory formation (Data S5). Previous studies demonstrate a role for OBPs in alcohol sensitivity and tolerance [108–113], and change expression in the MB following long-term memory formation [114]. Similarly, *Obp69a* expression decreases as a result of male flies being housed in groups [115]. We found that *Obp69a* expression was increased following alcohol exposure (Data S5), which is consistent with our observation that one day of intermittent ethanol-odor treatment decreases social behavior, mimicking previously demonstrated isolation phenotypes [37]. An upregulation of OBPs suggests a molecular signature through which alcohol can alter memory formation.

Another category of genes implicated in sensory responses include the large and diverse Cytochrome P450 family which encodes enzymes with a wide spectrum of monooxygenase and related activities. *Cyp4g1* was upregulated following one day of intermittent ethanol-odor exposure (Figure 3A) and *Cyp28d1* after three days of intermittent alcohol exposure (Figure 3C). *Cyp4g* genes catalyze the synthesis of cuticular hydrocarbons that serve a critical role in chemical communication and pheromonal response [116]. Pheromonal response is integral to social behavior such as courtship and aggression in insects [117–120]. A related gene *Cyp6a20* is regulated by social experience and consequently affects aggression [121–123]. Similarly, *Cyp4d21* is required for male courtship and mating success [124]. Our data suggest that ethanol induces molecular changes in chemosensory genes that have the potential to induce changes in social behavior, thus altering behavioral state.

### Alcohol increases gene expression associated with immune response

Our data demonstrates that genes involved in innate immune signaling have the most pronounced differential expression that occurs with repeated ethanol-odor pairing cycles (Figure 5, 6). Previous studies in humans with AUD, and in mammalian and fly models, have demonstrated similar gene expression changes, suggesting that the innate immune gene transcriptional response to alcohol exposure is involved in AUD pathobiology [125–132]. Furthermore, disruption of neuroimmune gene expression, or suppression of neuroimmune signaling pathways, reduces alcohol consumption [129, 133–139].

In mammals, toll-like receptors (TLR), are a key component of neuroimmune activation, and are widely thought to be a major contributor to alcohol dependence [140–142]. Pharmacological inhibition of TLR4 using naltrexone reduces ethanol preference in rodents [143, 144]. The *Drosophila* immune deficiency (Imd) pathway shares homologous components downstream of TAK1/TAB in the mammalian TLR pathway (Figure 6B) [145, 146]. We observed an upregulation of genes in both the Toll and Imd pathways over an extended three-day exposure cycle, as opposed to a single day of repeated alcohol exposure (Figures 5A, 6B, Data S5). This provides further evidence that alcohol exposure leads to a persistent aberrant innate immune signaling response, which persists even 48 hours after the final alcohol exposure.

The Toll and Imd pathways act together to regulate nuclear factor-κB, a critical factor in the innate immune response, and mediate differential expression of antimicrobial peptide encoding genes via distinct nuclear factor-κB-like transcription factors [145, 147–151]. An antimicrobial peptide, DptB, in *Drosophila* head fat body, was demonstrated to be important for long-term associative memory [152]. Furthermore, a number of differentially expressed Toll and Imd pathway genes have been previously implicated in alcohol sensitivity and tolerance, including *relish, spatzle, drosocin, serpins, Toll, Myd88, astray* and *Imd* [111, 132, 153–155].

Our data demonstrate that the two clusters of genes that are differentially expressed following repeated intermittent ethanol-odor exposure, but sustain these changes following 24 hr and 48 hr of ethanol deprivation, are heavily associated with innate immune signaling pathways (Figure 5B, 5C). A protein-protein network analysis using these genes revealed enrichment in key components and transcriptional targets of both the Toll signaling pathway including *Spz*, *Drosomycin*, *Defensin*, and *Metchnikowin*, and Imd signaling pathway including *PGRPs*, *Relish*, *Attacin, Cecropin, Diptercin, Drosocin*, and *Metchnikowin* (Figure 6). This shows intermittent ethanol-odor exposure, and subsequent ethanol deprivation, induces a sustained upregulation in innate immune response that is correlated with a profound shift in social behaviors.

### Social behavior and immune response

Social interactions are an important part of maintaining the overall health of a community. Although the mechanisms underlying social behavior are complex, recent findings indicate that immune signaling plays an important role in the regulation of social behaviors in a wide array of animals including insects, amphibians, and mammals [156]. Studies in rodents have demonstrated that the immune system is involved in anxiety behaviors [157–159]. Furthermore, in rodents, neuroimmune signaling influences both social behaviors, as well as spatial learning and memory [157, 160, 161]. Unravelling the link between behavior and neuroimmune signaling is challenging, we hypothesize that the increased social interaction in conjunction with increased innate immune gene expression that we observe is consistent with the notion that the neuroimmune signaling system regulates anxiety behaviors.

Interestingly, PGRPs, which are the receptors that sense pathogens, were recently shown to be required in octopamine neurons in order to behaviorally avoid pathogenic bacteria in *Drosophila* [162]. Octopamine neurons are required for a number of social behaviors including aggression and courtship in *Drosophila* [163], crickets [164], and ants [165]. Thus, ethanol-induced upregulation of these receptors could potentially alter the social behavior in response to pathogenic stimuli by affecting octopamine neuron function.

Another mechanistic link between immune signaling and social behavior that emerges from our study includes the Cytochrome P450 family of genes. As mentioned above, in insects these genes are associated with pheromonal response. Across phyla, from plants to insects and mammals, this family of genes is also associated with resistance to environmental stress like temperature and toxins [166–169]. In humans, heavy alcohol use leads to oxidative stress in the liver through increased metabolism via the cytochrome P450 system [170, 171]. We propose that the acute toxic properties of alcohol initiate changes in expression of Cytochrome P450 genes, that alter pheromone response and related social behavior. Further alcohol exposure and deprivation cycles then activate more prolonged immune responses, namely through Imd signaling. How these immune responses consequently affect social behavior remain to be characterized.

## Conclusion

*Drosophila* is a powerful model organism that provides novel insights into the underlying mechanisms of AUD. By studying the various facets of AUD, behavioral and neurobiological, we demonstrate that the repeated ethanol-odor pairing/deprivation cycles led to a distinct behavioral state characterized by an increase in social, including anxiety-like behaviors. We found increased neurotoxin and immune response gene expression associated with the increased social behaviors. We propose a model where alcohol disrupts the function of associative memory circuits required for stereotyped social behaviors by perturbing innate immune signaling.

Transcriptional regulation, particularly the immune signaling genes, plays an important role in the pathobiology of AUD. Further investigation of the roles of immune signaling genes, and their effects on specific regions of the brain and behavioral responses may lead to improved therapeutic strategies that can be used to identify and treat AUD. Gaining an increased understanding of AUD-related behavioral responses and gene expression changes in *Drosophila* to elucidate correlations between molecular signaling pathways, neural circuits, and behaviors with high specificity therefore has the potential to reduce gaps between preclinical disease models and clinical trials. Although the specific mechanisms involved in behavioral responses, the underlying neural circuits, and associated gene expression changes in response to alcohol remains to be elucidated, our study provides the framework to further investigate these relationships.

## Material and methods

### Fly Husbandry and maintenance

*Drosophila melanogoaster* wild-type (Berlin) flies were raised on cornmeal agar food at 24°C and 70% humidity with a 14:10 Light:Dark cycle. 3-5 day-old male flies were used throughout the entirety of this work. Male flies were collected 24 hours post eclosion, isolated under light CO_2_, given 2 days to recover, and then used for behavioral or molecular assays. Sibling age-matched control flies were trained simultaneously then tested either for alcohol-associated memory, spontaneous behavior in an open field arena, or frozen for transcriptomic analysis. Fly line listed in the Key Resources Table.

### Odor cue-induced ethanol memory

Sibling-matched and age-matched flies were trained and tested as previously described [11, 21]. Memory experiments were performed with 30 flies/vial, that underwent three spaced training sessions per day, for one or three days (Figure 1A). Training vials used were perforated 14 mL canonical culture tubes with mesh lids. In each training session, flies were presented with a session of 10 min of an unpaired odor, followed by 10 min of paired odor with ethanol, followed by a 50 min rest, before subsequent trainings. The odors were made up in mineral oil and used at a concentration of 1:36 odorant:mineral oil, with ethyl acetate or isoamyl alcohol as the odor. 10 mins of 90:60 (EtOH:Air) exposure caused a 6.6 ± 0.9mM (~0.03 g/dL) internal body ethanol concentration. Alcohol/odor exposure was administered in two distinct training paradigms: acute intermittent exposure, consisting of one day of three spaced trainings (Figure 1Ai), and chronic intermittent exposure, consisting of three days of three spaced trainings (Figure 1Aii, 1Aiii). To control for any inherent odor preferences all experiments were performed in reciprocal (averaged between alternative order of odors); a pair of reciprocal tests was used as biological n = 1. For the one day intermittent exposure groups, alcohol vapor was delivered to flies on 1% agar and supplemented with yeast pellets immediately after training. For the three day intermittent exposure groups, alcohol vapor was delivered to flies maintained on yeast-sugar (5% each) agar (1%).

Memory testing was performed in a Y-Maze (Figure 1B) where flies were given a choice between the paired and unpaired odor. Testing was done 24 hrs following the cessation of vapor treatment for the one day intermittent exposure group, and 24- or 48-hrs following vapor exposure for the three day intermittent exposure group. Conditioned preference index (CPI) was calculated by counting the number of flies that moved towards the paired odor, subtracting the flies that moved towards the unpaired odor, and dividing that number by the total number of flies.

Preference for the ethanol-paired odor was expressed as a conditioned preference index (CPI), where a positive CPI indicates preference and a negative CPI implies aversion to the ethanol-associated odor (Figure 1Bi, 1Bii, 1Biii). As an odor control, flies were trained in the same training paradigms with odors only, and these naïve flies had no significant preference for the odors used (Figure S1).

### Open Field Arena Behavior Experiments

Sibling and age-matched flies underwent three spaced training sessions per day, for one or three days (Figure 1A), following which flies were deprived of alcohol for 24 hrs or 48 hrs (Figure 1Ai, 1Aii, 1Aiii). Flies were then placed in the FlyBowl apparatus to record spontaneous behavior [53]. The FlyBowl apparatus is an open-field walking arena that accommodates 4 groups of 10 male flies in each of the 4 behavior arenas [53] (Figure 1C). In the FlyBowl, flies’ locomotion and social behaviors are observed using a software called FlyBowlDataCapture (FBDC) by video recording a group of freely behaving flies in the FlyBowl arena [172].

Each 30 min recording session consisted of 4 arenas, or bowls, containing 10 male wild-type Berlin flies each. The video files were then processed to generate trajectory data using a computer vision tracking software called FlyTracker [54]. JAABA, a behavioral annotating software, was then used to quantify specific behaviors, or classifiers, using the FlyTracker output files [55]. Each behavior (attempted copulation, back up, chaining, chase, crabwalk, jump, pivot center, pivot tail, righting, stops, touch, and walk) was then quantified across time for both the ethanol-paired odor and odor only groups (Figure S2, S3, S5).

### Behavior Tracking

We used automated tracking and behavior analysis algorithms to generate behavioral output data sets from the video recordings. A modified version of ctrax [172] called FlyTracker [54], was used to produce trajectory data compatible with JAABA, a behavioral annotating software. The benefits of using FlyTracker instead of Ctrax are that FlyTracker produces less tracking errors such as identity swaps and loss of fly identities, therefore eliminating the need to manually fix errors [35]. However, due to the bouts of clustering or chaining the groups of flies in the FlyBowl perform, identity swaps still occur, and sometimes there are loss of fly identities. Despite this, FlyTracker still corrects the number of fly tracks to preserve the original fly tracks (10 fly identity tracks in each video). Once the JAABA compatible files and per-frame folder is created from FlyTracker, the directories containing the video data and JAABA files are organized into “experimental directories” for JAABA use [55]. Following the creation of perframe features, behavior classifiers are assigned using JAABA, which is outlined below.

### Behavior Classification

Once the per-frame features are constructed in the perframe directory for all experimental directories, JAABA can use these directories to train subprograms, known as classifiers, which annotate one behavior. When adding experimental directories as training videos for a classifier, the JAABA program automatically checks if the added directories have the correct per-frame features the classifier is currently using. A classifier needs a foundation of training data to be able to annotate behavior correctly and automatically from videos. Once done training, the classifier can label its predictions for the user to test its performance on other videos. If there are incorrect predictions, the user can re-train the classifier. To verify the accuracy of the classifier to correct label behavior, we used JAABA’s program called Groundtruthing Mode. We used multiple classifiers from a preexisting list known as BABAM classifiers [36], for the exception of the chaining classifier, which we created. The chaining classifier corresponds to the observation of a group of 3 or more flies facing the same direction, where each fly inside of the chain is touching the tail of another fly or the flies at the ends of the chain have flies touching their tails.

To generate the behavioral data from each classifier, we used JAABA’s program, JAABADetect, which generates a score .mat file per classifier. To produce this as a batch process line, we wrote a MATLAB program that gathers the list of experiment folders containing the JAABA trajectory data as well as the perframe features, and then creates score files with an inputted list of classifiers. We extracted only the binary behavioral data from the score .mat files and wrote each fly’s identity track data to a csv file for one behavior to make this data parsable in R. We wrote an R script to parse the csv data from each experiment to produce time graphs containing data points of 15 second bins, graphing an average number of occurrences of each behavior across these 15 seconds. To get the 15 sec bins of data for an experiment, we first transformed each individual fly’s data in the csv file for behavior from frames of data to 0.5 sec intervals of data. In these 0.5 sec intervals, rounded the mean of the intervals to get the binary behavior of a 0.5 sec data point. Once all the fly individual’s data was transformed, we get the experiment’s number of occurrences in 0.5 sec intervals by summing up all fly’s data for each 0.5 interval. We used the subset of experiments for a behavior time graph and for each of these experiments, and transformed the 0.5 sec intervals to 15 sec intervals by summing up the number of occurrences in the 15 sec intervals (30, 0.5 intervals go into a 15 sec bin). Finally, to obtain the average number of occurrences (per experiment), we calculated the mean of all experiment’s 15 sec intervals and plotted the smooth shaded areas with standard deviation. We then applied a smoothing function to get the mean of that point and the next two points and update the point with the mean. The total sum of each behavioral occurrence per fly for each classifier was then calculated. The average of the total sums of the 10 flies per bowl was then used to transform the data into z-scores.

### Neural Circuit Identification

To investigate the potential circuits involved in the behavioral outputs identified in each condition, the total sum of the behavioral occurrences in the paired ethanol-odor group was compared to the odor only control group. For example, attempted copulation decreased 24 hours following one day of intermittent ethanol-odor exposure, and increased 24 and 48 hours after three days of intermittent ethanol-odor exposure, when compared to their respective odor only control group. The directionality of each behavior compared to its control was used to identify circuits using the BABAM software. For example, if the ethanol + odor pair group had a greater total number of occurrences of a given behavior, the directionality would be specified as “increased” on the BABAM software. Once the behavioral classifier and direction are specified, the brain regions associated with the behavioral trends were projected onto a map of the *Drosophila* brain using a colormap to signify the relative p-value associated with the brain region. The dark red represents the lowest p-value (p<0.0001) and the dark blue represents the highest significant p-value (p=0.05). Regions that were not significantly implicated in the behavioral shifts remain black on the brain projections. Only the brain regions involved in all social behaviors and all locomotor behaviors in each condition were then used to narrow down the search for regions involved in the behavioral trends.

### RNA-seq Data Collection

RNA-seq data were collected from whole heads of sibling, and age-matched flies trained in the odor cue-induced ethanol training paradigm, following which flies were deprived of ethanol for 24 hrs or 48 hrs (Fig. 1Ai, 1Aii, 1Aiii, 1D). Biological replicates were performed for odors only (n = 4) and ethanol-odor trained (n = 4) conditions, which included reciprocal odor groups. For each replicate, 30 whole fly heads were collected by flash freezing flies using liquid nitrogen, followed by sieve separation. RNA was extracted from frozen heads, and RNA libraries were prepared using NuGEN’s Encore Complete RNA-Seq kit, and RNA sequencing was done using Illumina NextSeq 550.

### RNA-seq Data Analysis

Analysis of the RNA-seq data was done using Trajectory Inference and Mechanism Exploration with Omics data in R (TIMEOR) [57]. Overall, the time-series RNA-seq and multi-omics analysis method TIMEOR was run in the command line and through the web-interface (via RShiny) [174] to process all replicates for all three timepoints, automatically cluster genes based on inferred gene trajectory dynamics, and produce temporal and per timepoint gene ontology (GO) analysis results. Specifically, the raw data uploaded into TIMEOR and sequenced reads were run through FastQC [175] with default parameters, to check the quality of raw sequence data and filter out any sequences flagged for poor quality. No sequences were flagged. Sequenced reads were then mapped to release 6 *Drosophila melanogaster* genome (dm6) [176] using both Bowtie2 [177] and HISAT2 [178]. Across all replicates, Bowtie2 returned a higher median percent total alignment, and were converted to sorted bam files (*bowtie2 -x dm6_genome -1 replicate_R1_001.fastq.gz -2 replicate_R2_001.fastq.gz -S out.sam > stout.txt 2> alignment_info.txt; samtools view -bS out.sam > out.bam; rm -rf out.sam; samtools sort out.bam -o out.sorted.bam*). Each aligned replicate .bam file was then converted to read counts per gene using HTSeq [179] to produce fragments per kilobase of transcript per million mapped reads (FPKM) counts. All resulting replicate files were merged to create a matrix of FPKM expression counts for each gene within each replicate. Using principal component analysis to visualize several normalization and correction methods, Trimmed-Mean of M-values normalization and Harman [180] correction helped remove poor quality replicates. Correlation plots between replicates revealed the highest correlation between replicates. Within TIMEOR, DESeq2 [173] using the Wald test of significance was chosen to perform differential expression analysis by first estimating size factors and dispersion, followed by negative binomial gaussian linear model fitting and the Wald test of significance.

We leveraged TIMEOR’s automatic unsupervised clustering [56] to group genes with similar gene expression trajectories across all conditions. We compiled a list of differentially expressed genes (using the less conservative p<0.05 cut-off) from each condition (Table S7). The resulting 430 differentially expressed genes were automatically clustered into nine groups representing similar gene dynamic trajectories through TIMEOR. The clusters were generated using TIMEOR’s Euclidean distance between genes and Ward D2 [181] method between clusters. For each cluster and within each timepoint, TIMEOR performed GO analysis using clusterProfiler [182] to search for enriched terms within biological processes. GO terms that are significantly enriched for a group of genes are depicted as dot plots. *Dot plots* depict enriched GO terms (x-axis) vs. the ratio of genes that are enriched for that GO term (y-axis). Each dot radius shows the enriched gene count for that GO term and the dot color indicates the GO term significance (using the adjusted p-value). TIMEOR performed these steps automatically, producing intermediate and publication ready figures at each step. Software and algorithms used here are listed in the Key Resources Table.

### Statistical Analysis

#### Behavior statistics

The behavioral datasets were assessed with parametric statistics using SPSS. A Levene’s test was used to determine homogeneity of variance and Shapiro-Wilk test was used to assess normality of the data prior to running parametric statistics. For all experimental raw data, the mean and standard error were calculated and shown. The behavioral data was evaluated using a repeated measures ANOVA to compare the ethanol-paired odor and odor only groups across time, * p < 0.05, **p < 0.01, ***p < 0.001. For comparisons of behavioral trends across the different conditions, a 2-way ANOVA was conducted, * p < 0.05, **p < 0.01, ***p < 0.001.

#### Neural circuit identification statistics

To investigate the potential circuits involved in the behavioral outputs identified in each condition, the total sum of the behavioral occurrences in the ethanol-odor group was compared to the odor only group. The directionality of each behavior compared to it’s control helps to narrow down the circuits identified with the BABAM software. Once the behavioral classifier and direction are specified, the brain regions associated with the behavioral trends are projected onto a map of the Drosophila brain using a colormap to signify the relative p-value associated with the brain region. The dark red represents the lowest p-value (p<0.0001). Only the brain regions involved in all social behaviors and all locomotor behaviors in each condition were then used to narrow down the search for regions involved in the behavioral trends.

#### RNA-seq statistics

Differential gene expression analyses were performed on 30 whole fly heads after the period of forced deprivation. We used the time-series RNA-seq and multi-omics analysis method, within TIMEOR, to generate, analyze, and visualize gene expression data. We used R to create volcano plots to display fold change vs p-value statistics between odor control and odor trained libraries. Enriched GO terms for genes with a p-value < 0.05 were plotted on a dot plot, where GO processes with the largest gene ratios are plotted in order of gene ratio, using TIMEOR. The size of the dots represents the number of genes associated with the GO term, and the color of the dot represents the adjusted p-values.

## Acknowledgements

Heartfelt thanks go to Kristin Branson (Janelia Research Campus), Alice Robie (Janelia Research Campus), and Mayank Kabra (Janelia Research Campus) for assistance with FlyBowl, ctrax and JAABA, and to members of Fly Olympiad (Janelia Research Campus) for assistance with FlyBowl experiments. Thanks also to Jason Ritt (Carney Institute, Brown University) for advice on analysis of behavior data. This work could not have been done without the assistance of Serge Picard (Janelia Research Campus) and the Janelia Research Campus NeuroSeq Team. Thank you also to Fred Davis (Janelia Research Campus) and Emily Petrucelli (Southern Illinois University Edwardsville) and the Brown University Center for Computational Molecular Biology (CCMB) for preliminary analysis of the RNA-Seq data. Special thanks to Galit Shohat Ophir (Bar Ilan University) for assistance with data collection for RNA-Seq and advice on the manuscript. And finally, thanks to John McGeary (Brown University) and members of the Kaun Lab for advice on data visualization, data interpretation and writing.

## Author Contributions

N.M.D performed RNA-Seq data visualization and interpretation, writing and editing. K.S.M performed behavior data visualization and interpretation, circuit data generation using BABAM datasets, statistical analysis of behavior data, writing and editing. A.C. developed the time-series RNA-seq and multi-omics analysis method, performed RNA-seq Data analysis, and assisted during behavioral data analysis strategizing. T.B. performed behavior data analysis and visualization. R.A. provided original ideas and generated behavior data. U.H. provided original ideas, mentoring and funding support. E.L. provided mentoring and funding support for A.C. K.R.K. provided original ideas, generated RNA-Seq data and behavior data, and provided mentoring, funding support, data interpretation, writing and editing.

