## Supplementary material for "Neuromolecular and behavioral effects of ethanol deprivation in *Drosophila*": Figures S1 - S6

**Table of Contents** **page**

Title page.................................................. 1
Supplemental Figure 1.................................................. 2
Odor control data
Supplemental Figure 2.................................................. 3
Behavioral time graphs 24 hours following one day of ethanol exposure
Supplemental Figure 3.................................................. 5
Behavioral time graphs 24 hours following three days of ethanol exposure
Supplemental Figure 4.................................................. 7
Locomotor behavioral trends are associated with distinct brain regions
Supplemental Figure 5.................................................. 9
Behavioral time graphs 48 hours following three days of ethanol exposure
Supplemental Figure 6.................................................. 11
Gene expression changes with alcohol exposure and forced abstinence

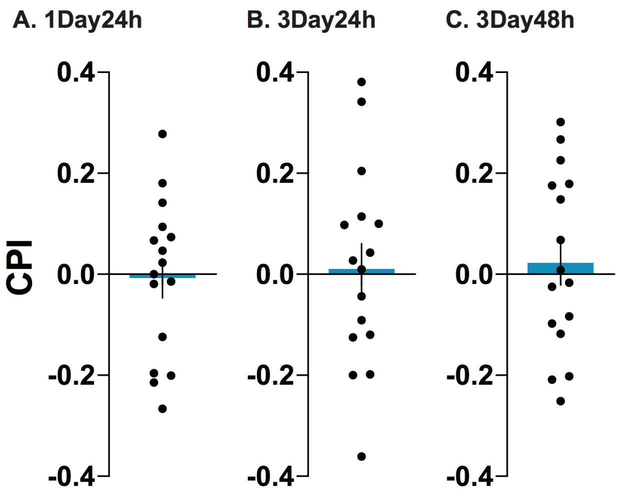


**Figure S1. Odor control data**

(A – C) As an odor control, flies were trained in the same training paradigms described in Figure 1A with odors only, and these naïve flies had no significant preference for the odors used.

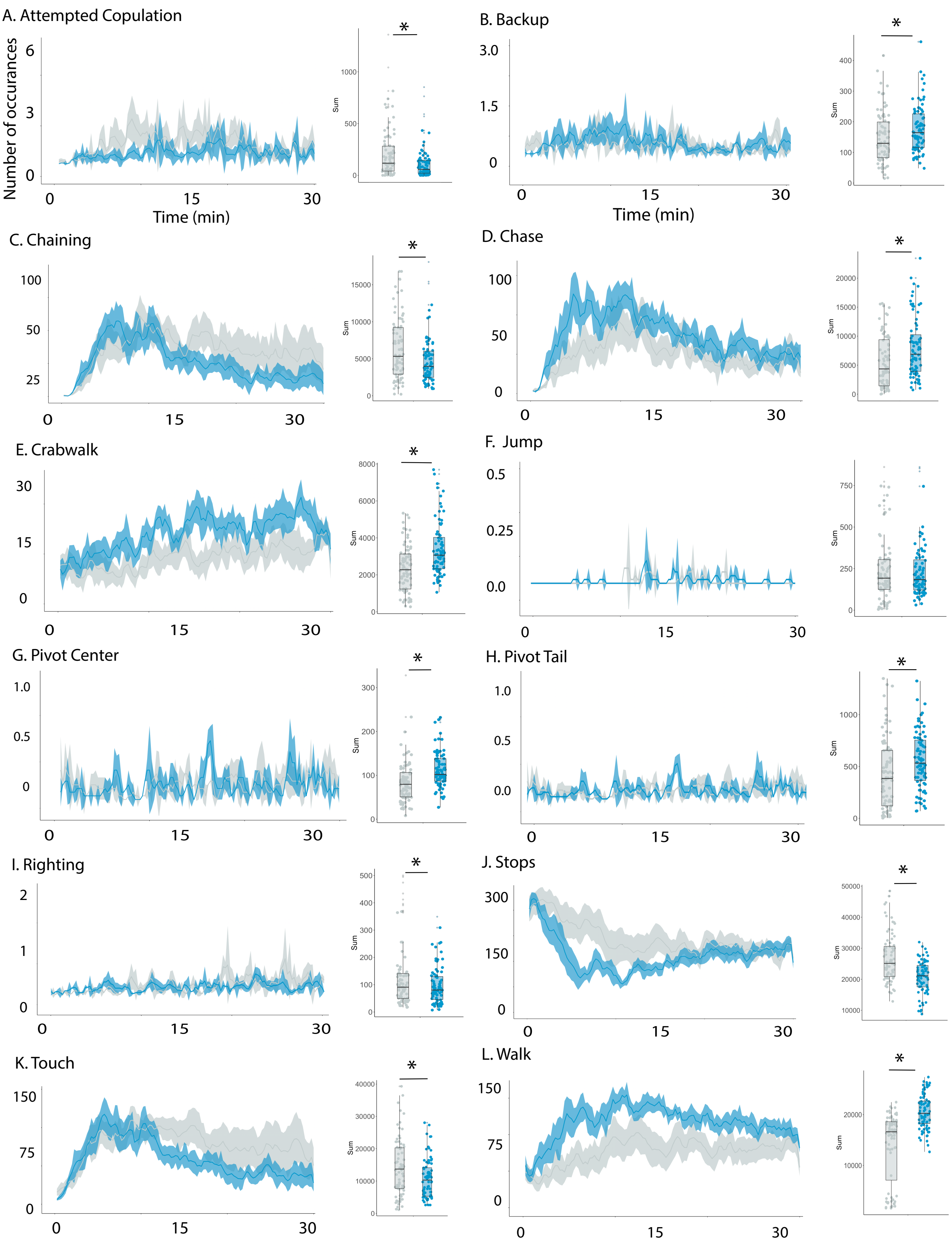
A. Attempted Copulation
Number of occurances
Time (min)
B. Backup
Time (min)
C. Chaining
D. Chase
E. Crabwalk
F. Jump
G. Pivot Center
H. Pivot Tail
I. Righting
J. Stops
K. Touch
L. Walk
Sum
*

**Figure S2. Behavioral time graphs 24 hours following one day of ethanol exposure**

The average number of occurrences of each behavioral classifier across the 30 minute video with the ethanol group in blue and the odor control group in gray. The total sum of each behavior per bowl is shown in the box plot to the right of each time graph, *$p<0.05$. (A) attempted copulation ($p = 0.007$), (B) backup ($p = 0.02$), (C) chaining ($p = 0.01$), (D) chase ($p = 0.002$), (E) crabwalk ($p < 0.001$), (F) jump ($p = 0.8$), (G) pivot center ($p < 0.001$), (H) pivot tail ($p = 0.009$), (I) righting ($p = 0.04$), (J) stops ($p < 0.001$), (K) touching ($p < 0.001$), (L) walk ($p < 0.001$).

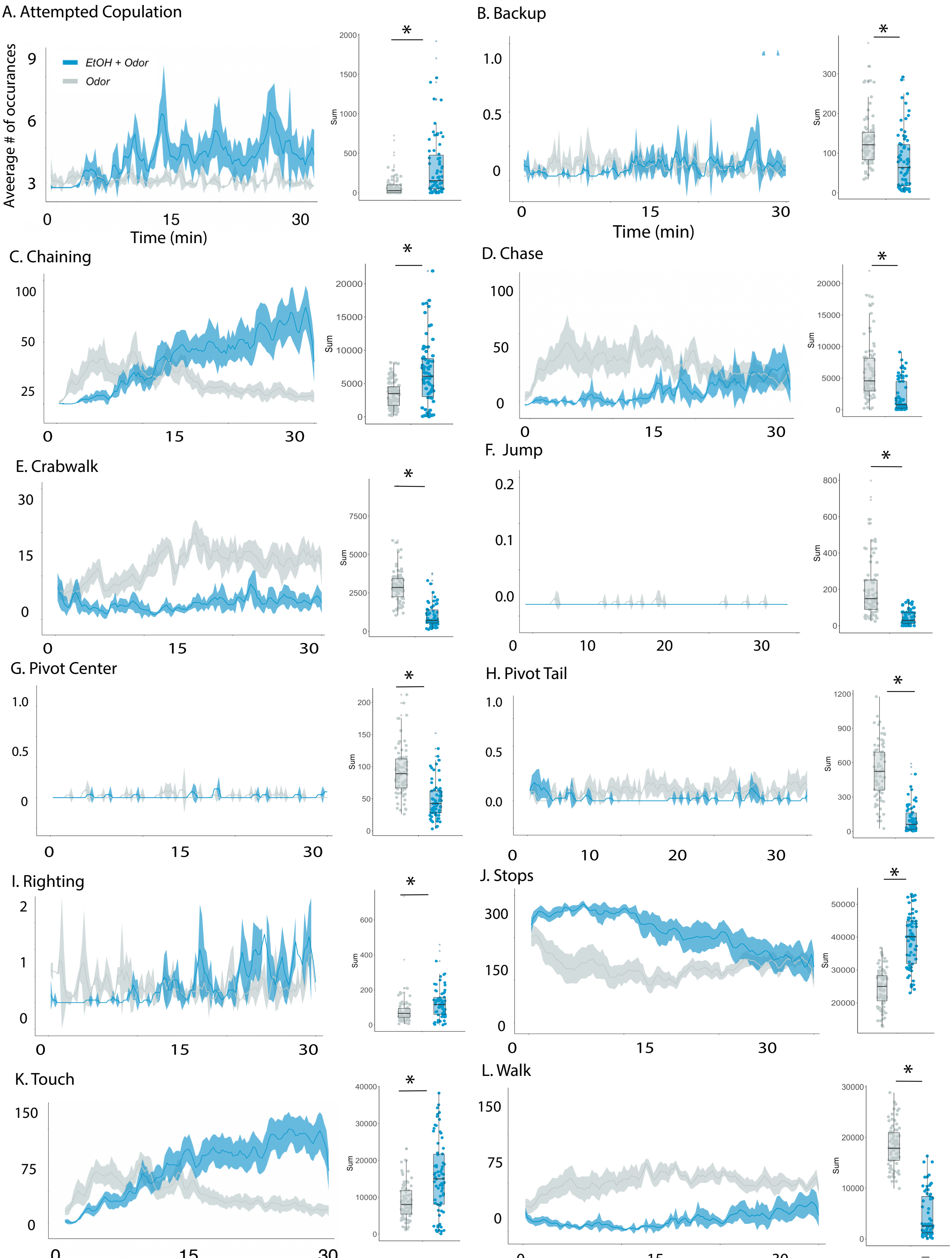
A. Attempted Copulation
EtOH + Odor
Odor
Average # of occurances
Time (min)
Sum
*
B. Backup
C. Chaining
D. Chase
E. Crabwalk
F. Jump
G. Pivot Center
H. Pivot Tail
I. Righting
J. Stops
K. Touch
L. Walk

**Figure S3. Behavioral time graphs 24 hours following three days of ethanol exposure**

The average number of occurrences of each behavioral classifier across the 30 minute video with the ethanol group in blue and the odor control group in gray. The total sum of each behavior per bowl is shown in the box plot to the right of each time graph, *$p<0.05$. (A) attempted copulation ($p < 0.001$), (B) backup ($p < 0.001$), (C) chaining ($p < 0.001$), (D) chase ($p < 0.001$), (E) crabwalk ($p < 0.001$), (F) jump ($p < 0.001$), (G) pivot center ($p < 0.001$), (H) pivot tail ($p < 0.001$), (I) righting ($p = 0.009$), (J) stops ($p < 0.001$), (K) touching ($p < 0.001$), (L) walk ($p < 0.001$).

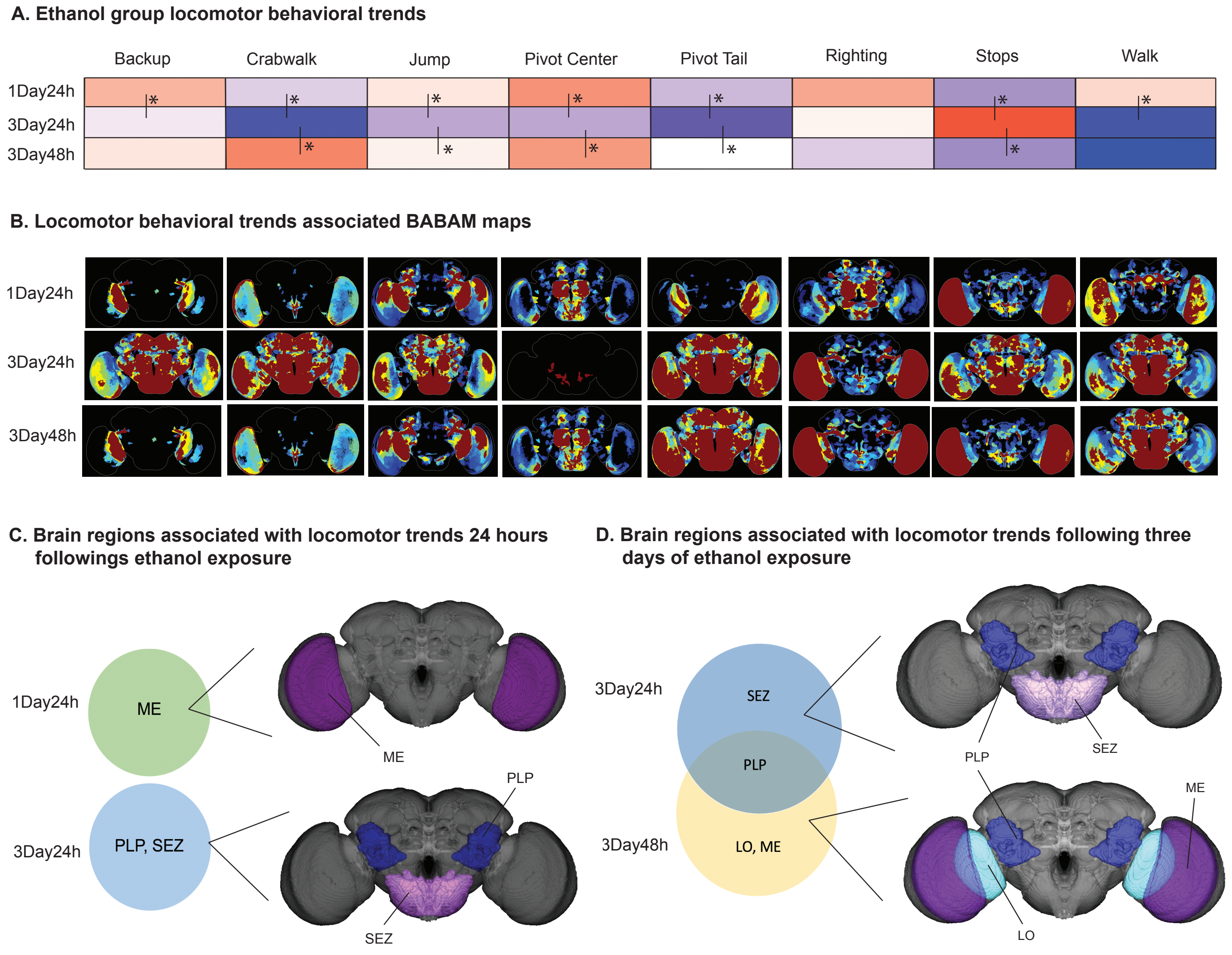


**Figure S4. Locomotor behavioral trends are associated with distinct brain regions**

(A) Locomotor behavioral trends associated with 24 hours of forced abstinence after 1 day (1Day24h) or three days (3Day24h) of ethanol exposure or 48 hours of forced abstinence after three days of ethanol exposure (3Days48h). The heatmap shows the EtOH group behavior occurrences sums compared to the odor controls total sums. Red indicates an increase in total sum and blue indicates a decrease in total sum. Statistical significance was evaluated by ANOVA, post hoc Tukey, to compare EtOH trends across conditions, $*p<0.05$. (B) BABAM projections of the brain regions implicated in the behavioral trends found for each behavioral feature. Dark red indicated the lowest p-value, blue indicates the highest p-value. (C) A Venn diagram showing the brain regions implicated in the locomotor behavioral trends found in each 24 hour forced abstinence conditions, 1Day24h and 3Day24h. The medulla (ME) was unique to 24 hours of forced abstinence following 1 day of exposure and the posterior lateral protocerebrum (PLP) and subesophageal zone (SEZ) were unique to 24 hours of forced abstinence following 3 days of exposure. The brain regions in each section of the Venn diagram are projected onto the brain maps to the right of the Venn diagram. (D) A Venn diagram showing the brain regions implicated in the locomotor behavioral trends found in each three-day ethanol exposure conditions, 3Day24h and 3Day48h. The medulla (ME) and lobula (LO) was unique to 48 hours of forced abstinence, the subesophageal zone (SEZ) was unique to 24 hours of forced abstinence and the posterior lateral protocerebrum (PLP) was common among both groups. The brain regions in each section of the Venn diagram are projected onto the brain maps to the right of the Venn diagram.

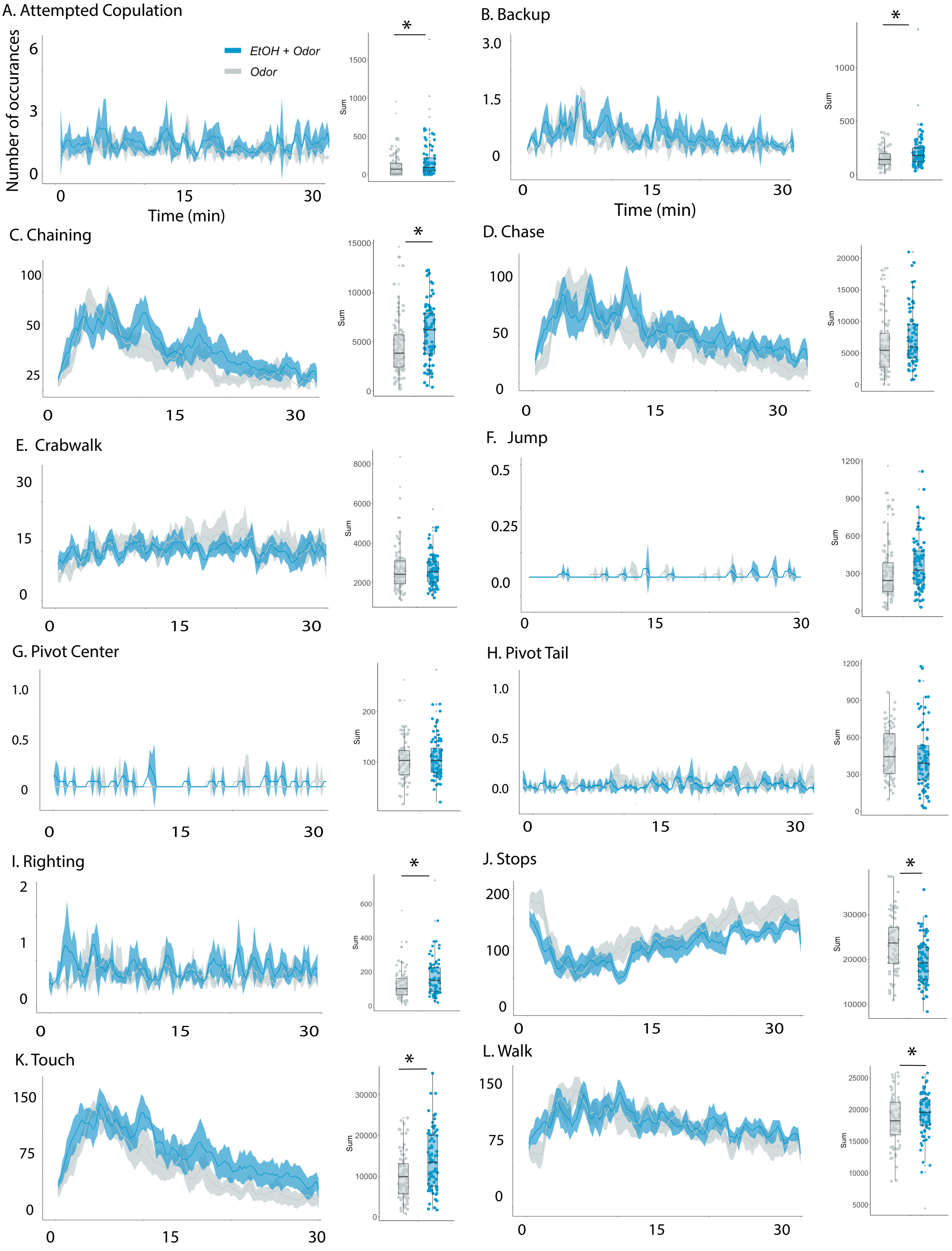

A. Attempted Copulation
EtOH + Odor
Odor
Number of occurances
Time (min)
Sum
*
B. Backup
C. Chaining
D. Chase
E. Crabwalk
F. Jump
G. Pivot Center
H. Pivot Tail
I. Righting
J. Stops
K. Touch
L. Walk

**Figure S5. Behavioral time graphs 48 hours following three days of ethanol exposure**

The average number of occurrences of each behavioral classifier across the 30 minute video with the ethanol group in blue and the odor control group in gray. The total sum of each behavior per bowl is shown in the box plot to the right of each time graph, *$p<0.05$. (A) attempted copulation ($p = 0.03$), (B) backup ($p = 0.004$), (C) chaining ($p = 0.003$), (D) chase ($p = 0.07$), (E) crabwalk ($p = 0.7$), (F) jump ($p = 0.1$), (G) pivot center ($p = 0.3$), (H) pivot tail ($p = 0.3$), (I) righting ($p = 0.003$), (J) stops ($p < 0.001$), (K) touching ($p < 0.001$), (L) walk ($p < 0.001$).

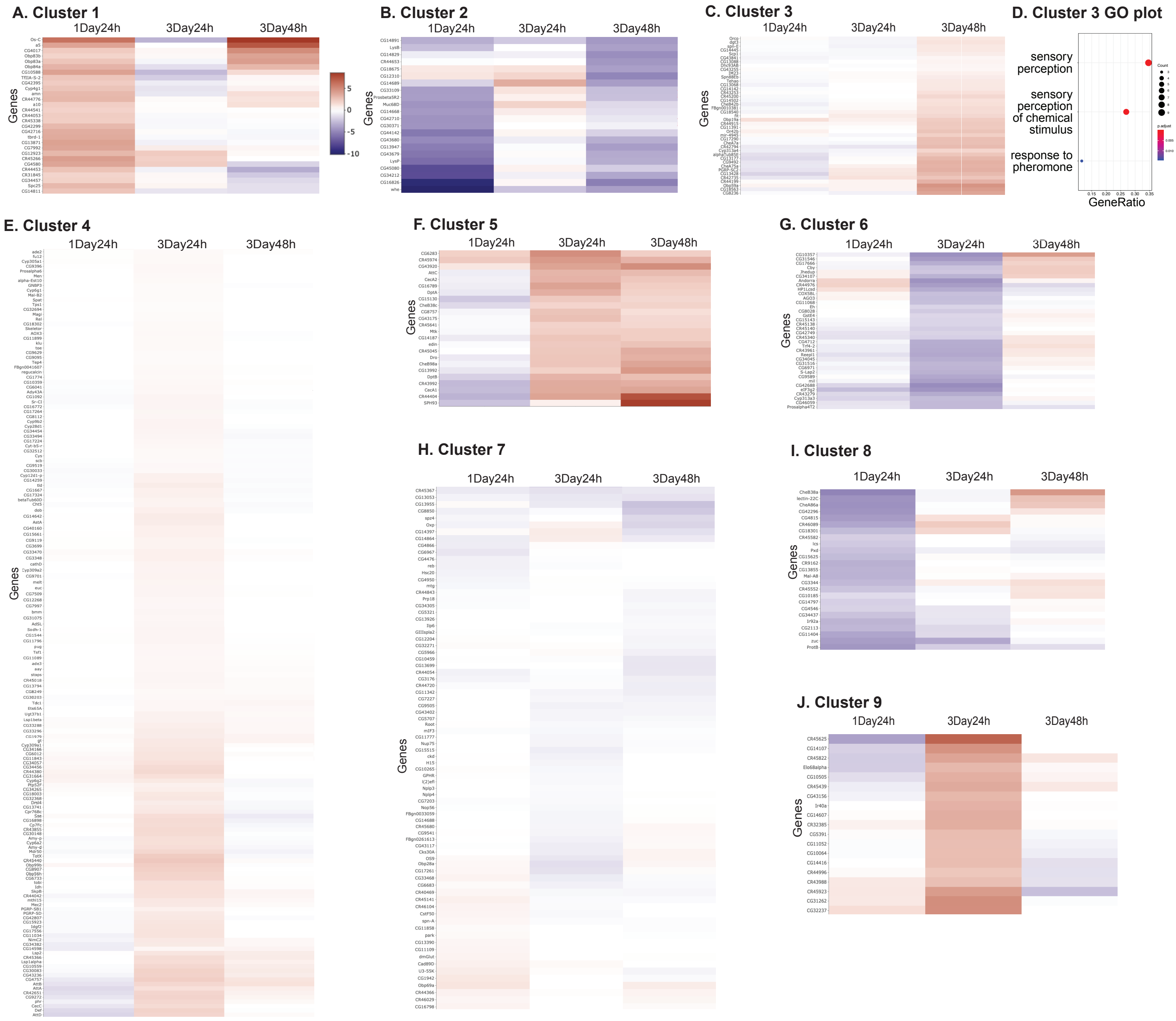


**Figure S6. Gene expression changes with alcohol exposure and forced abstinence**

(A - C; E - J) We used a clustermap to identify clustering of differentially expressed genes, with a p-value < 0.05, from each condition, 24 hours after one day of spaced intermittent training (1Day24h), 24 hours after three days of spaced intermittent training (3Day24h), and 48 hours after three days of spaced intermittent training (3Day48h), resulting in 9 clusters representing similar gene dynamic trajectories. (D) GO enrichment analysis on genes within cluster 3, plotted on a dot plot. The 3 GO processes with the largest gene ratios are plotted in order of gene ratio. The size of the dots represents the number of genes associated with the GO term, and the color of the dot represents the p-adjusted values.
